# T1234: A distortion-matched structural scan solution to misregistration of high resolution fMRI data

**DOI:** 10.1101/2024.09.19.613939

**Authors:** Chung (Kenny) Kan, Rüdiger Stirnberg, Marcela Montequin, Omer Faruk Gulban, A Tyler Morgan, Peter Bandettini, Laurentius (Renzo) Huber

## Abstract

**Purpose:** High-resolution fMRI at 7T is challenged by suboptimal alignment quality between functional data and structural scans. This study aims to develop a rapid acquisition method that provides distortion-matched, artifact-mitigated structural reference data.

**Methods:** We introduce an efficient sequence protocol termed T1234, which offers adjustable distortions. This approach involves a T1-weighted 2-inversion 3D-EPI sequence with four spatial encoding directions optimized for high-resolution fMRI. A forward Bloch model was used for T1 quantification and protocol optimization. Twenty participants were scanned at 7T using both structural and functional protocols to evaluate the utility of T1234.

**Results:** Results from two protocols are presented. A fast distortion-free protocol reliably produced whole-brain segmentations at 0.8mm isotropic resolution within 3:00-3:40 minutes. It demonstrates robustness across sessions, participants, and three different 7T SIEMENS scanners. For a protocol with geometric distortions that matched functional data, T1234 facilitates layer-specific fMRI signal analysis with enhanced laminar precision.

**Conclusion:** This structural mapping approach enables precise registration with fMRI data. T1234 has been successfully implemented, validated, and tested, and is now available to users at our center and at over 50 centers worldwide.

## 1.) Introduction

High-resolution fMRI promises to reveal fine-scale neural information flow across layers and columns. However, common approaches of high-resolution fMRI with Cartesian EPI are limited by EPI phase artifacts dominantly caused from eddy-current induced trajectory imperfections and B_0_-related geometrical distortions. A survey by the ISMRM brain function study group found that the most limiting factor of the field of high-resolution fMRI is image registration^1^. This is despite the fact that researchers commonly invest significant time (≈10 min) in acquiring high-quality anatomical data with MP2RAGE^2^.

In this study, we build on previous works^3-12^ by developing, implementing, characterizing and validating a novel acquisition method: T1234, involving T_1_-weighted acquisition with 2 inversions using a 3-dimensional EPI readout in 4 directions. This sequence provides structural reference images with high anatomical contrast, adjustable geometrical distortions, and mitigated artifacts in 3-4 minutes.

### 1.1) Historical heritage of T1-EPI

Since the advent of EPI-based fMRI^13-14^, geometric distortions have been a recognized challenge, and have subsequently been addressed through dewarping techniques^15^. T1-weighted EPI protocols were introduced around the same time^16-18^. With 7T fMRI at higher resolution, Huber et al^3^ used T1-weighted EPI in first attempts to estimate voxels of upper and deeper layers in the distorted functional space of constraint fMRI slabs. High-resolution T1-EPI was then combined with more advanced strategies, including slice-wise inversion time cycling for broader 2D-EPI coverage^5-6^, simultaneous multi-slice readouts^9^, 3D-EPI, and segmented readouts^11,8^. Van der Zwaag et al. named their approach T123, referring to T1-imaging with two 3D-EPIs. Further developments have combined advanced 3D-EPI techniques with alternative MT-preparation methods^4,10^. These advancements, achieving sub-millimeter resolutions on 7T scanners, were driven by the goal of facilitating mesoscale layer-fMRI analyses. However, none of these approaches has yet gained widespread adoption. We suspect that this is largely due to the inherent noise in EPI, which typically demands long acquisition times and averaging. Moreover, some noise sources stem from consistent EPI trajectory imperfections, which cannot be mitigated through averaging alone.

Our work here builds on the method by Stirnberg et al^8^. We further optimize this technique by integrating multi-phase encoding directions^10^, interleaved dual-polarity readouts^19-20^, CNR optimizations by means of TI-specific flip angles that are larger than in conventional Look-Locker approaches, and we combined it with a forward Bloch model^16^ for T1 quantification. This sequence is tested for two protocols. A fast distortion-free whole brain protocol that can serve as a layer-fMRI optimized alternative to conventional MP2RAGE. And a distortion-matched protocol that allows layer segmentation in the functional data without spatial warping.

## 2.) Methods

We tested the T1234 sequence in 20 MRI sessions across three 7T scanners: two SIEMENS Terra scanners and one SIEMENS 7T Plus scanner. MRI data were acquired under the NIH-IRB (93-M-0170, ClinicalTrials.gov: NCT00001360).

### 2.1 Acquisition

Key protocol parameters included: 0.8mm isotropic resolution, Skipped-CAIPI 1×3z1, segmentation factor of 14^7-8^, matrix size of 232×232×186, and a phase partial Fourier factor of 6/8. We utilized a relatively high segmentation factor to minimize distortions and T2*-blurring. For a complete list of parameters, see: https://github.com/layerfMRI/Sequence_Github/tree/master/T1234. The loop structure of this sequence^8,21^ is illustrated in Figure 1 for the proposed protocol.

**Fig. 1:**
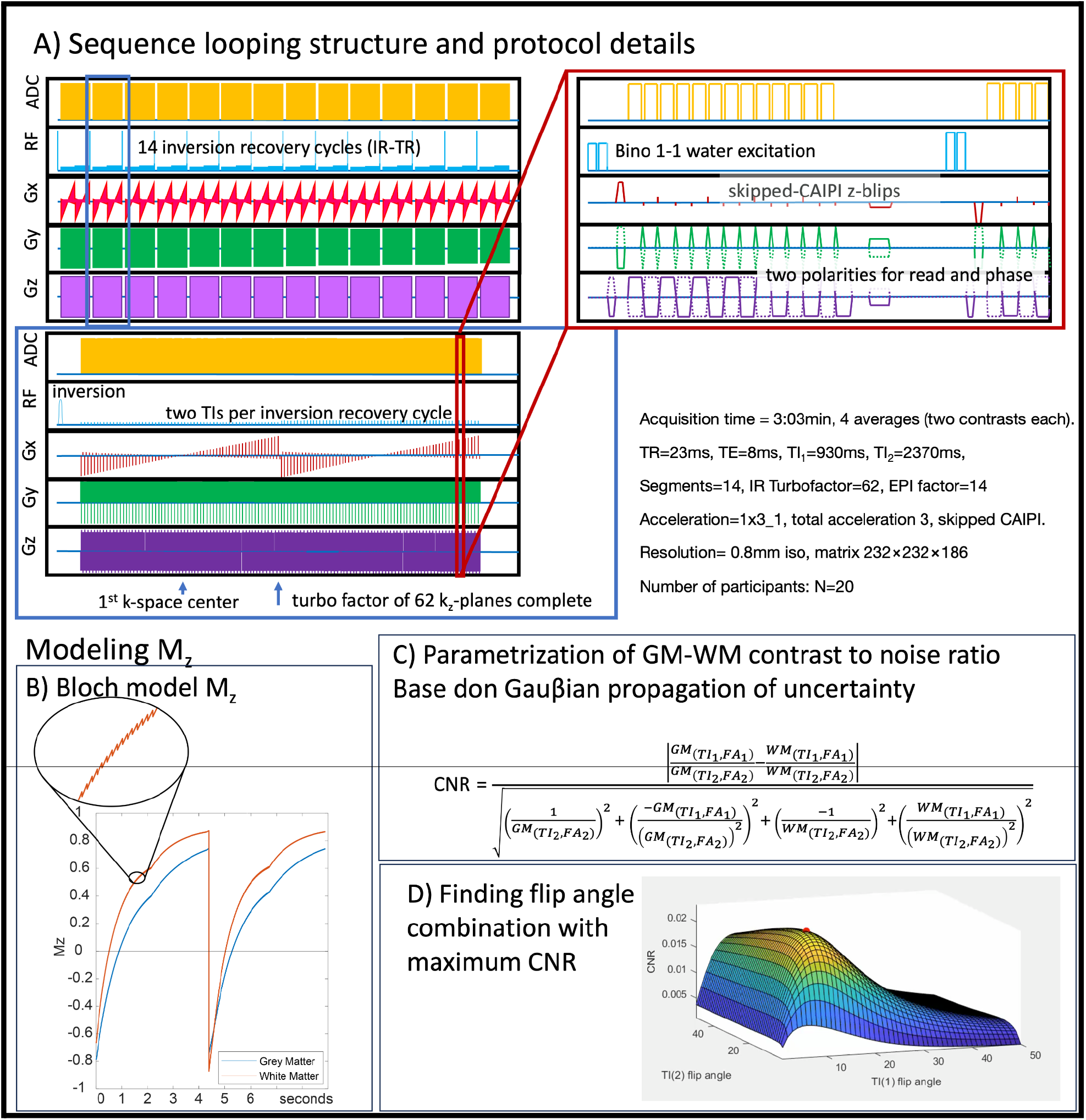
Sequence diagram and corresponding expected magnetization. **A)** Individual panels are zoomed-in sections of one complete volume acquisition of a given read and phase polarity combination. This illustrates the sequence looping structure across interleaved multi-shot segments, inversion recovery cycles, kz planes, and ky lines. This entire cycle of a volume acquisition is repeated four times, incorporating all combinations of reversed read and phase directions. **B)** This panel schematically depicts the Bloch model that represents the expected Mz magnetization in the inversion-recovery cycle with non-selective excitation pulses. The saw-tooth pattern in the zoomed-in section results from the segment-specific excitation flip angle. The two distinct apparent relaxation segments correspond to two different flip angles for the first and second inversion times. **C)** Based on this Bloch model, the expected gray matter-white matter contrast-to-noise ratio (CNR) can be parameterized as a Gauβian propagation of uncertainty. **D)** This panel shows the CNR as a function of the two flip angles for the first and second inversion times, respectively. The local maximum is highlighted with a red dot.

A single volume of a bias-field corrected T1-weighted signal was reconstructed from pairs of inversion times (comparable to the UNI-image in MP2RAGE, Equation 3, of the article by Marques et al.^2^). In addition to the increased SNR achieved through averaging, the following corrections were implemented:

1. The use of two read directions helps to mitigate typical EPI low spatial frequency artifacts, known as “Fuzzy Ripples,” which result from imperfections in k-space trajectories and off-resonance effects^19,12^.
2. Two images acquired in reverse phase encoding directions have opposite distortions. The adjustable segmentation level and phase bandwidth in T1234 can be matched to the effective echo spacing of the functional data, thereby aligning distortions at the acquisition level. Alternatively, with distortions in opposite directions, researchers can estimate the distortion field and retrospectively synthesize any desired level of geometric warping, similar to the topup method^22^, here implemented in AFNI.

In five participants, in addition to the T1234 acquisition, we also tested the acquisition using a single phase encoding direction while keeping the effective echo spacing identical to that of a conventional fMRI protocol for 7T at standard resolutions^23^.

Additionally, we conducted four functional experiments to assess the usability of this tool for functional T1 mapping. These included three 12-minute runs of checkerboard stimulation with four inversion times (TIs) and IR-TRs (Figure 1A) of 3.8 seconds, and a 40-second TR with two TIs.

### 2.2 Reconstruction and Analysis

Complex-valued averaging was performed after parallel image reconstruction and coil combination using IcePat^24^. Nyquist ghost correction was carried out using the vendor’s standard two-parameter implementation^25,-27^. The same navigator and GRAPPA reference data were used for both EPI read polarities. For each phase encoding direction polarity, a different set of phase navigators and GRAPPA reference data was acquired and used.

Motion correction of complex-valued image space data was performed using ANTS^28^. Layers were extracted using the Layer-Toolbox framework (Barilari: https://github.com/marcobarilari/layerfMRI-toolbox) with LayNii v2.6.0^29^, FreeSurfer^30^ v7.4.1, and PreSurfer 1f5ad28 as a wrapper for SPM12 tools^31^.

### 2.3 Analysis of T1 estimation

To predict the optimal distribution of variable flip angles across the inversion-recovery evolution, based on the employed loop structure (Figure 1), we implemented a forward Bloch simulation. This simulation allowed us to determine the excitation flip angles that provide the maximum contrast-to-noise ratio (CNR) between gray matter and white matter (Figure 2). The Bloch solver was also used to generate a lookup table (Figure 3A) that converts relative signal changes across inversion times to physical units of T1 in milliseconds. Given that the protocol used here employs relatively large flip angles (>10°), a simple Look-Locker analysis approach that assumes small flip angels is not justified. The spin-history of each readout alters the T1-relaxation curve depending on local B1 inhomogeneity-dependent excitation flip angles. As a first-order approximation of locally dependent Mz-saturation, we utilized a universal low-resolution template B1 map, previously obtained across the cortex. Simulation code and analysis scripts are available at: https://github.com/layerfMRI/repository/tree/master/T1234.

**Fig. 2:**
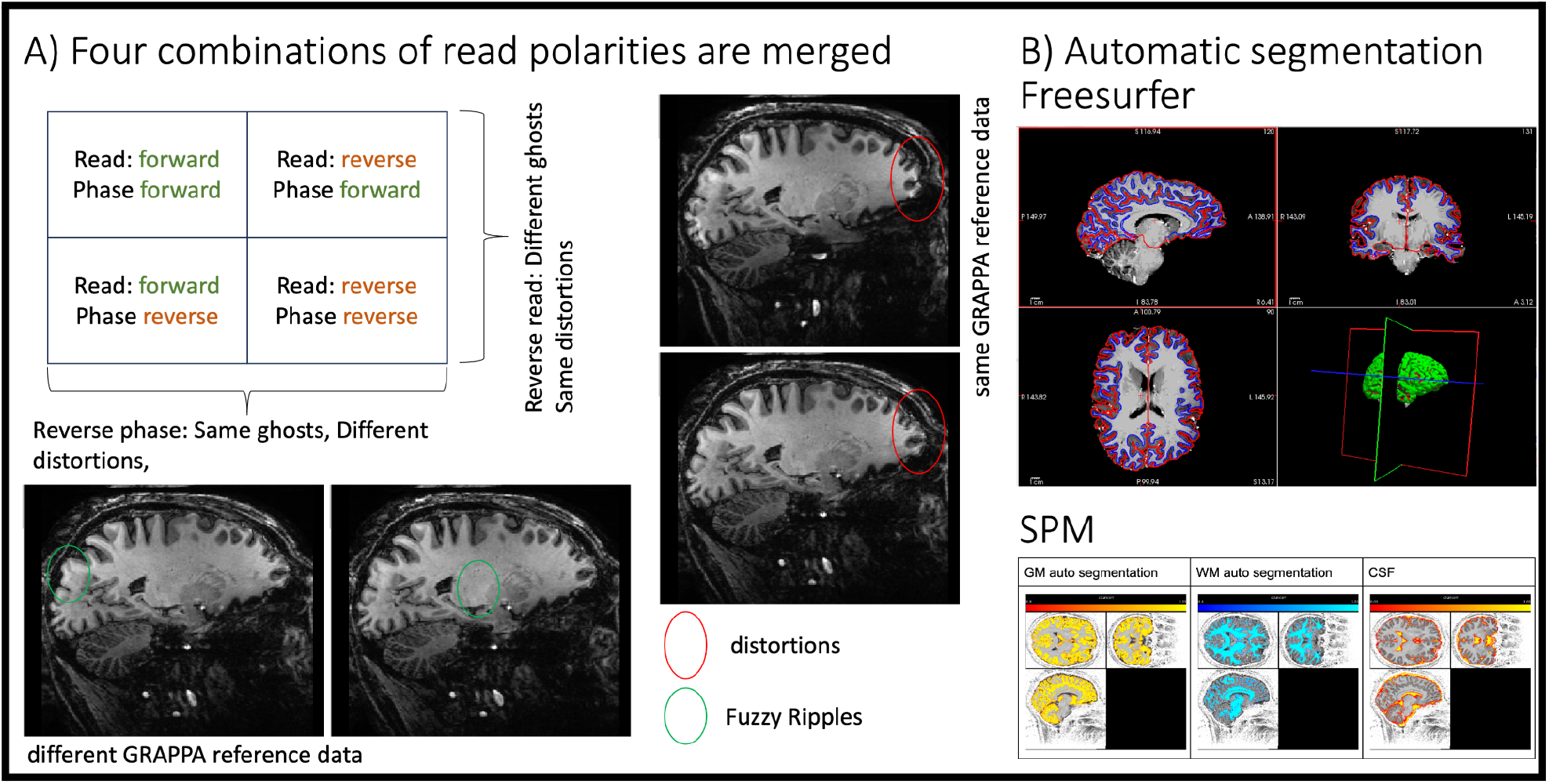
Merging four combinations of read and phase polarity imaging for tissue-type segmentation. **A)** Example image showing artifacts that are mitigated by combining read and phase polarities. The ellipses highlight artifact levels that are benignly inverted by reversing read and phase polarity. Green ellipses indicate “fuzzy ripples”, a low spatial frequency artifact caused by gradient trajectory imperfections at EPI ramp-sampling corners. For an animated version of this figure that more clearly depicts the fuzzy ripples, see here: https://github.com/layerfMRI/repository/blob/master/T1234/animation_2.gif **B-C)** After combining the four images, the data can be processed through mainstream automatic segmentation pipelines. Representative results from FreeSurfer and SPM are shown, respectively.

**Fig. 3:**
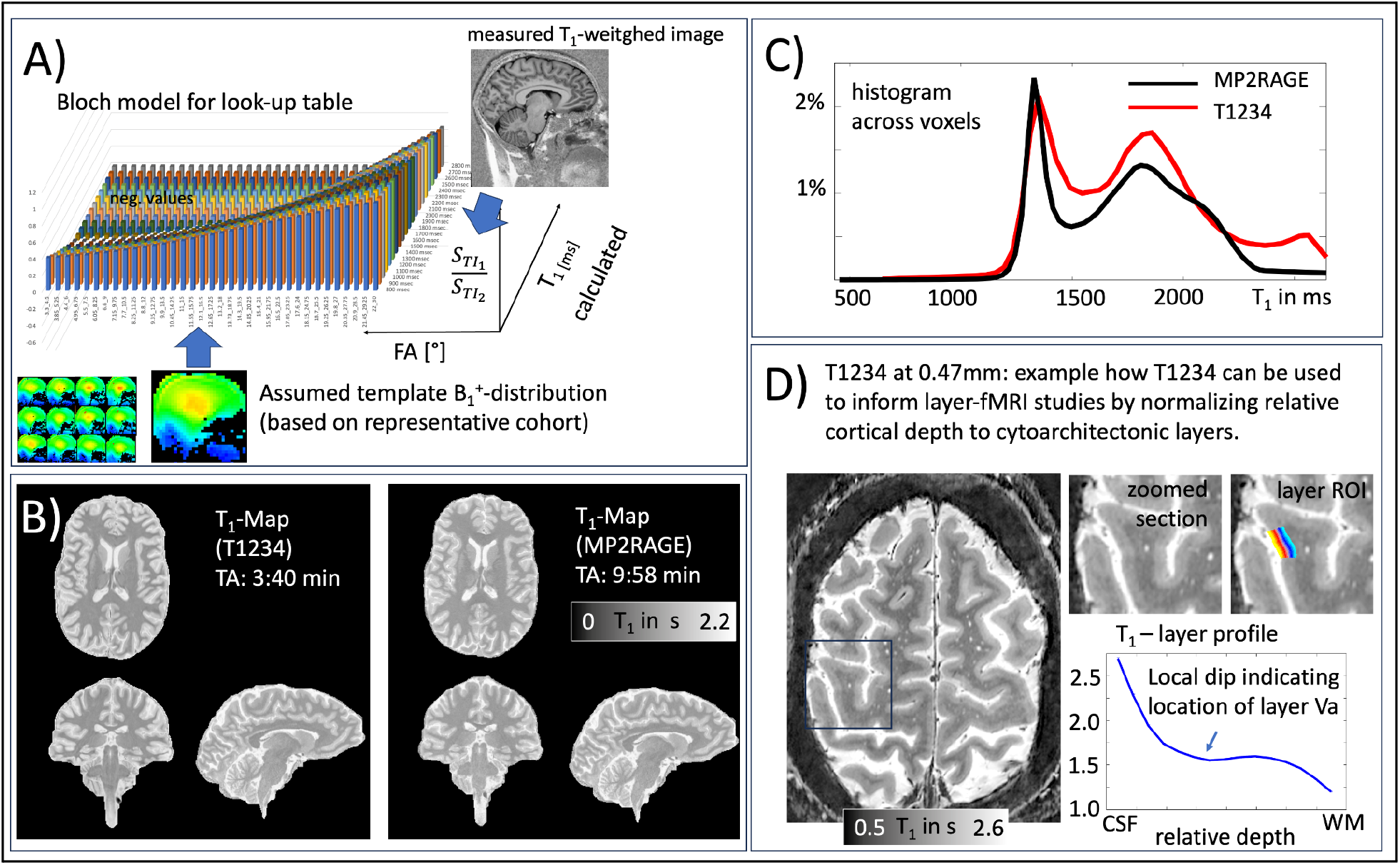
T1 quantification for the T1234 approach. **A)** The Bloch solver (Figure 1B) is used to generate a lookup table of expected relative MR signal intensities for T1 tissue types at both inversion times, depending on TR, FA, and the IR looping structure. **B-C)** Representative T1 map and corresponding histogram generated using the proposed approach, compared to a T1 map generated with MP2RAGE. The GM and WM peaks are largely identical. **D)** Example application of slab-selective T1234 for layer-fMRI. At higher spatial resolutions for layer-specific mapping, T1234 can be used to identify myeloarchitectonic landmarks to calibrate cortical depth to layers.

## 3.) Results

Figure 2 depicts representative results from one participant. The signal maps from the first inversion time show strong gray matter-white matter (GM-WM) contrast and good signal-to-noise ratio (SNR) with minimal graininess. However, these images also display conventional EPI artifacts, including geometric distortions and “fuzzy ripples” (Figure 2A). By combining data from multiple read and phase polarity directions, these artifacts can be mitigated, making the images suitable for standard tissue segmentation (Figure 2B).

Fig. 3A illustrates how T1234 data can be used to estimate quantitative T1-values. This is done by using a lookup table derived from a forward Bloch model to convert measured signal ratios of TI1 and TI2 into T1 values in milliseconds. The resulting T1 values are compared with those derived from MP2RAGE (a standard in the field) as maps (Figure 3B) and as histograms (Figure 3C). Slight deviations might arise due to unmodeled MT effects during the readout in either sequence, potentially incomplete inversion efficiency, imperfections in the MP2RAGE T1 model for relatively short TRs (4.5s), or inaccuracies in flip angle estimation in the T1234 Bloch model based on the universal B1 distribution approach used here.

Quantitative T1 mapping can be valuable for calibrating geometric cortical layers^32^ to myelo-architectonic layers^33^, using the shape of quantitative T1 profiles as anatomical landmarks. This is exemplified in Figure 3D with a modified slab-selective high-resolution T1234 protocol at 0.47mm isotropic.

We found that T1234 is highly robust across participants, sessions, scanners (Figure 4A), and runs (Figure 4B).

**Fig. 4:**
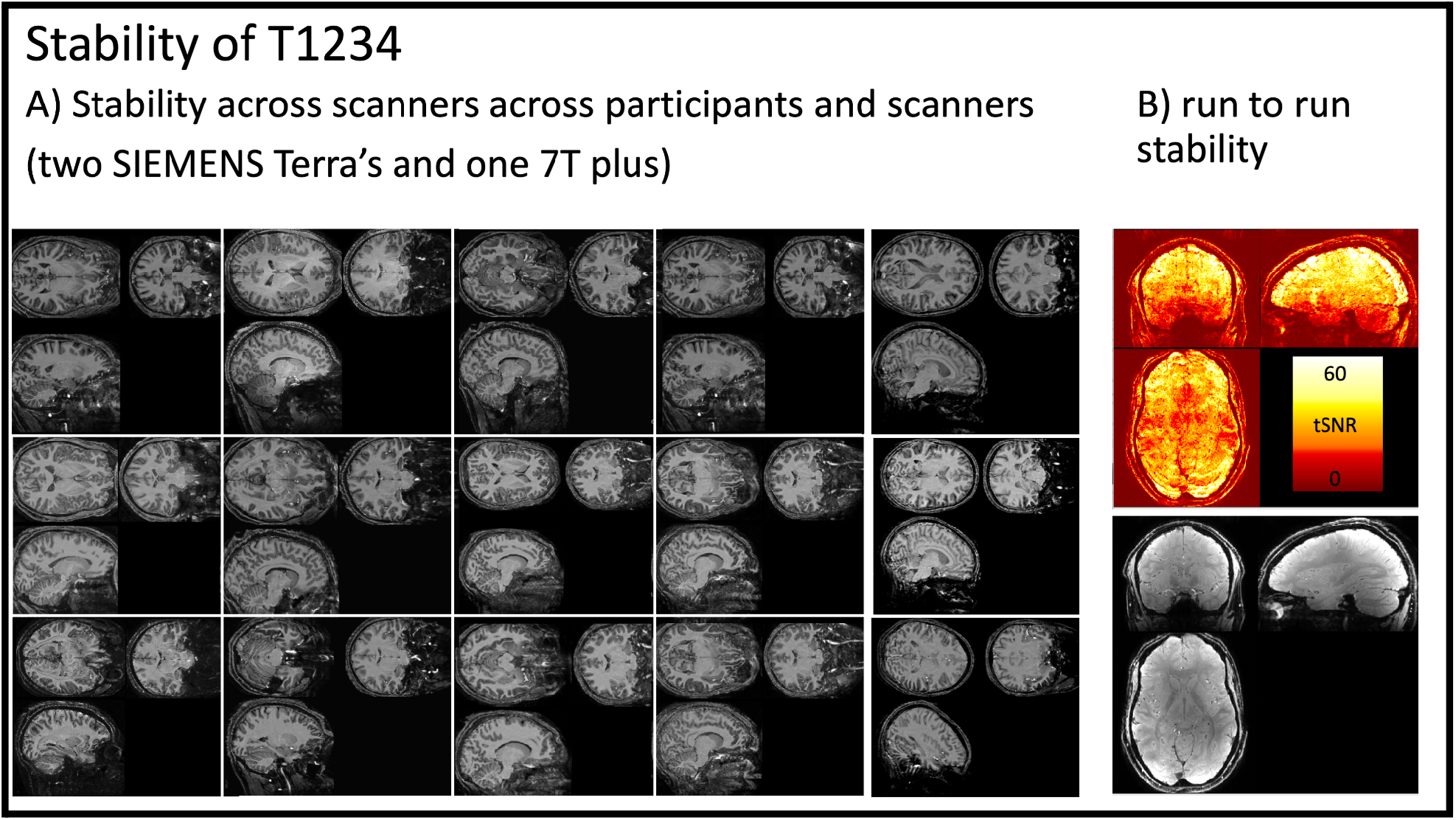
Stability of T1234. **Panel A)** depicts T1234 maps across 15 scan sessions, showing consistency across participants and different scanner models. The protocol reliably provides strong T1 contrast. **Panel B)** shows run-to-run stability of the protocol in the form of tSNR maps, calculated as the mean divided by the standard deviation.

To demonstrate the value of the distortion-matched T1234 sequence for layer-fMRI research, we compared laminar fMRI signal pooling using T1234-derived tissue borders with signal pooling using conventionally derived MP2RAGE borders. Fig. 5 shows that laminar signal pooling from distortion-matched T1234-derived layer segmentation retains spatial signal modulation across cortical depth with higher spatial fidelity.

**Fig. 5:**
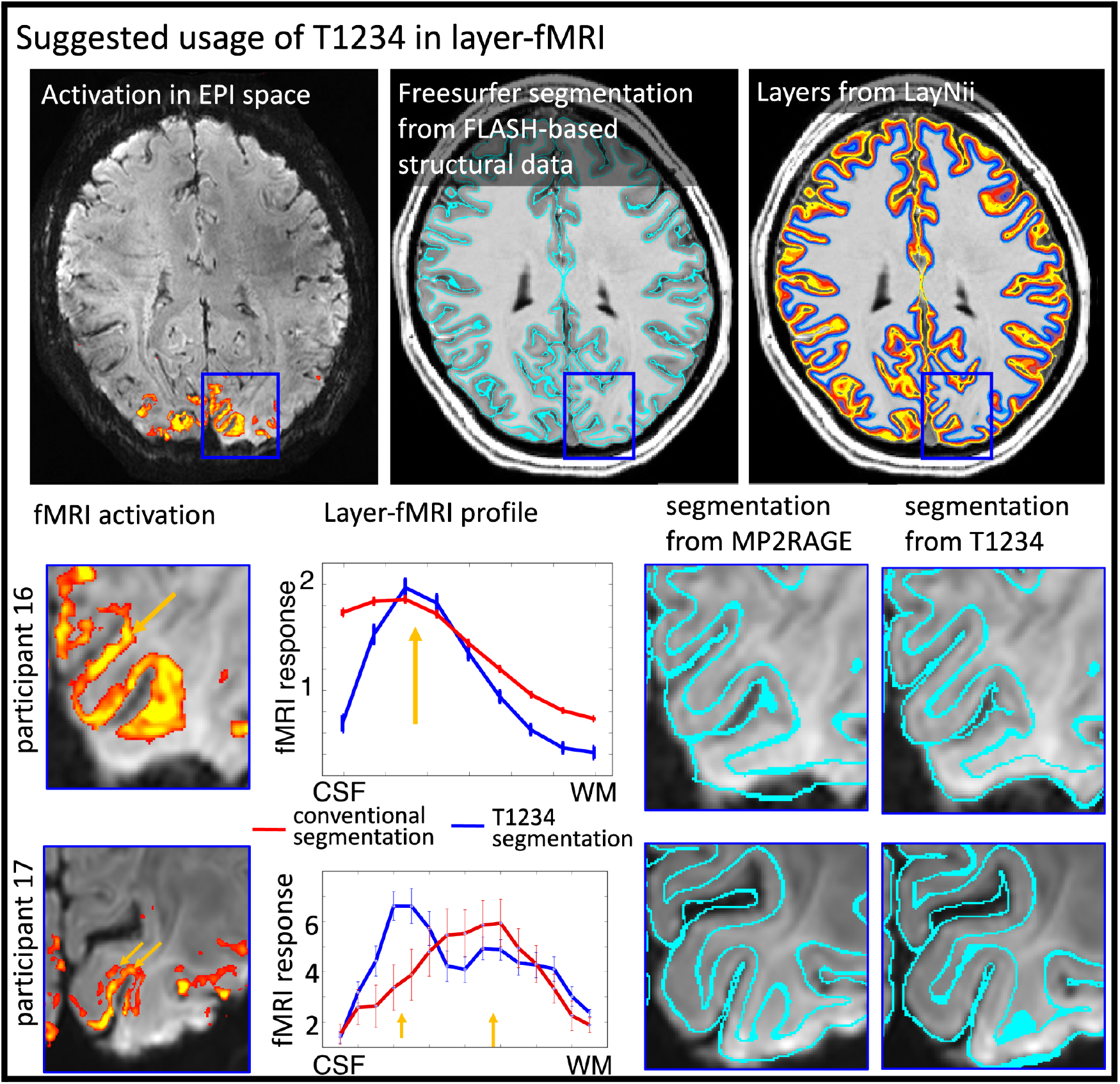
Proposed usage of T1234 for high resolution layer-fMRI applications Conventional layer-fMRI activation mapping typically uses FLASH-based structural reference data. These structural reference data are segmented in software tools like FreeSurfer to estimate layers for the extraction of activation scores across cortical depth. However, if the alignment between tissue-type borders (as derived from structural reference data) does not perfectly match the locations of functional activation data, the resulting layer profiles are compromised. The panels for participants 16 and 17 illustrate two challenges of the conventional layer-mapping approach. The orange arrows point to layer-specific activation patterns that follow the cortical ribbon at unique cortical depths. However, when these signals are pooled from compromised segmentation borders of conventional structural reference data, the depth-dependent peaks are blurred. When segmentation borders have higher spatial precision from T1234 data, the peaks of depth-dependent activations become visible in layer profiles.

## 4) Discussion

### 4.1) Importance of high-precision tissue type segmentation

Current high-resolution fMRI studies are often constrained by the spatial accuracy of tissue segmentation. Previous research has reported that manual correction of automatic and semi-automatic segmentation approaches require extensive labor^34,35^. This bottleneck in high-resolution fMRI has been recognized by the ISMRM Brain Function Study Group as the most significant challenge in mesoscale fMRI^1^. Our T1234 method has the potential to overcome this challenge by achieving high precision without the residual constraints and artifacts associated with other EPI-based structural mapping protocols.

### 4.2) Future work of T1234

#### 4.2.1 Optimizing this general T1234 protocol for special cases

Our T1234 approach is optimized for applications in layer-fMRI and whole-brain coverage. While this protocol facilitates straightforward use of FreeSurfer pipelines, it may be less efficient for applications focused solely on individual brain areas. For such cases, optimizing the acquisition protocol for higher resolutions and smaller field-of-views (FOV) prescriptions, as demonstrated in Fig 3D, might be more efficient.

#### 4.2.3 Limitations of universal B1 calibration

The protocol used here enables quantitative T1 estimation in physical units of milliseconds, assuming a universal B1+ pattern. This method has been effective for B1+ shimming in previous studies^36^, but it may not fully account for participant-specific local deviations from the representative mean. While this approach is sufficient for anatomical landmark identification, as demonstrated in Figure 3D, applications requiring more precise T1 quantification might benefit from acquiring a participant-specific B1 map. This can typically be done in under a minute.

#### 4.2.3 Limitations of EPI

The high segmentation factor proposed in this protocol allows for relatively short echo train lengths of 14, with an image echo time of 8ms. Although this is significantly shorter than conventional EPI readouts, it remains longer than those used in conventional FLASH-based structural protocols. As a result, the T1234 approach is more susceptible to fast signal decay induced by susceptibility gradients, particularly in brain regions with significant B0 inhomogeneity near air cavities.

## 5) Summary and conclusion

EPI-based acquisition of structural reference data has the potential to address registration challenges in high-resolution fMRI studies. However, except for VASO studies, where functional scans double as structural scans, such approaches have not gained widespread adoption. This is largely due to the long acquisition times and EPI artifacts associated with functional scanning, which diminish the appeal of using these methods for structural reference. In this study, we build on previous EPI protocols for structural imaging and propose a multi-directional EPI readout approach that addresses two critical limitations: distortion and artifact levels. Our protocol offers a fast, whole-brain scan in just 3-4 minutes and has proven to be robust for tissue type segmentation across different participants and scanners. Additionally, it has been found to improve cortical depth estimation in layer-fMRI studies. This sequence is available for sharing on 7T SIEMENS scanners and is currently being utilized by over 50 sites worldwide.

## 6) Acknowledgements

## Funding

This work was is supported by the NIH Intramural Program of NIMH/NINDS (#ZIC MH002884). MRI scanning was performed in the FMRIF core.

## Conflict of interest

Omer Faruk Gulban is an employee of Brain Innovation (Maastricht, NL). The work presented here may be partly specific to industrial design choices of SIEMENS Healthineers’ UHF scanners. This vendor is used in 83% of all human layer-fMRI papers (source: www.layerfmri.com/papers).

## Ethics

MRI data were acquired under the NIH-IRB (93-M-0170, ClinicalTrials.gov: NCT00001360). We thank Shruti Japee for guidance and support with respect to getting privileges for checking pregnancy tests and IRB.

## Support

We thank Hoan Le and Vinai Roopchansingh for help with computing hardware used for the simulation work conducted here.

## 7) Data Availability Statement

- Data of all participants can be found on Zenodo as two parts: https://doi.org/10.5281/zenodo.13366784 and https://doi.org/10.5281/zenodo.13376563
- Analysis scripts are available here: https://github.com/layerfMRI/repository/tree/master/T1234
- Scan protocols are available here: https://github.com/layerfMRI/Sequence_Github/tree/master/T1234
- The T1234 sequence with on-scanner reconstruction of coil-specific dual-polarity combination can be accessed via the Siemens C2P exchange platform: For European IP addresses https://webclient.eu.api.teamplay.siemens-healthineers.com/c2p and for US IP addresses https://webclient.us.api.teamplay.siemens-healthineers.com/c2p. Search for 3D-EPI VASO (by Stirnberg and Huber) provided by DZNE.

## 8) Diversity statement

Recent work in several fields of science has identified a bias in citation practices such that papers from women and other minorities are under-cited relative to the number of such papers in the field^37^. In the human layer-fMRI community the average of the gender citation bias is 84% male, 16% female (https://layerfmri.com/papers/). We obtained the gender of the first author of each reference. By this measure (and excluding self-citations to all authors of our current paper), our references contain 83% male first and 17% female first. This method is limited in that: (i) names, pronouns, and social media profiles used to construct the databases may not, in every case, be indicative of gender identity, and (ii) it cannot account for intersex, non-binary, or transgender people. We look forward to future work that could help us to better understand how to support equitable practices in science.

## Notes

https://doi.org/10.5281/zenodo.13366784

https://doi.org/10.5281/zenodo.13376563

